# Microbiome differential abundance methods produce disturbingly different results across 38 datasets

**DOI:** 10.1101/2021.05.10.443486

**Authors:** Jacob T. Nearing, Gavin M. Douglas, Molly Hayes, Jocelyn MacDonald, Dhwani Desai, Nicole Allward, Casey M. A. Jones, Robyn Wright, Akhilesh Dhanani, André M. Comeau, Morgan G. I. Langille

**Affiliations:** Department of Microbiology and Immunology, Dalhousie University; Department of Mathematics and Statistics, Dalhousie University; Department of Computer Science, Dalhousie University; Integrated Microbiome Resource, Dalhousie University; Department of Civil and Resource Engineering, Dalhousie University; Department of Pharmacology, Dalhousie University, Halifax, Nova Scotia, Canada

**Author notes:** These authors contributed equally (Jacob T. Nearing, Gavin M. Douglas).

## Abstract

Identifying differentially abundant microbes is a common goal of microbiome studies. Multiple methods have been applied for this purpose, which are largely used interchangeably in the literature. Although it has been observed that these tools can produce different results, there have been very few large-scale comparisons to describe the scale and significance of these differences. In addition, it is challenging for microbiome researchers to know which differential abundance tools are appropriate for their study and how these tools compare to one another. Here, we have investigated these questions by analyzing 38 16S rRNA gene datasets with two sample groups for differential abundance testing. We tested for differences in amplicon sequence variants and operational taxonomic units (referred to as ASVs for simplicity) between these groups with 14 commonly used differential abundance tools. Our findings confirmed that these tools identified drastically different numbers and sets of significant ASVs, however, for many tools the number of features identified correlated with aspects of the tested study data, such as sample size, sequencing depth, and effect size of community differences. We also found that the ASVs identified by each method were dependent on whether the abundance tables were prevalence-filtered before testing. ALDEx2 and ANCOM produced the most consistent results across studies and agreed best with the intersect of results from different approaches. In contrast, several methods, such as LEfSe, limma voom, and edgeR, produced inconsistent results and in some cases were unable to control the false discovery rate. In addition to these observations, we were unable to find supporting evidence for a recent recommendation that limma voom, corncob, and DESeq2 are more reliable overall compared with other methods. Although ALDEx2 and ANCOM are two promising conservative methods, we argue that those researchers requiring more sensitive methods should use a consensus approach based on multiple differential abundance methods to help ensure robust biological interpretations.

## Introduction

Microbial communities are frequently characterized with DNA sequencing. Marker gene sequencing, such as 16S rRNA gene sequencing, is the most common form of microbiome profiling and enables the relative abundances of taxa to be compared across different samples. A frequent and seemingly basic question to investigate with this type of data is: which taxa significantly differ in abundance between sample groupings? Newcomers to the microbiome field may be surprised to learn that there is little consensus on how best to approach this question. Indeed, there are numerous ongoing debates regarding the best practices for differential abundance (DA) testing with microbiome data (Allaband et al., 2019; Pollock et al., 2018).

One area of disagreement is whether read count tables should be rarefied (i.e., subsampled) to correct for differing read depths across samples (Weiss et al., 2017). This approach has been heavily criticized because excluding data could reduce statistical power and introduce biases. In particular, using rarefied count tables for standard tests, such as the t-test and Wilcoxon test, can result in unacceptably high false positive rates (McMurdie and Holmes, 2014). Nonetheless, microbiome data is still frequently rarefied because it can simplify analyses, particularly for methods that do not control for variation in read depth across samples. For example, LEfSe (Segata et al., 2011) is a popular method for identifying differentially abundant taxa that first converts read counts to percentages. Accordingly, read count tables are often rarefied before being input into this tool so that variation in sample read depth does not bias analyses. Without addressing the variation in depth across samples by some approach, the richness can drastically differ between samples due to read depth alone.

A related question to whether data should be rarefied is whether rare taxa should be filtered out. This question arises in many high-throughput datasets, where the burden of correcting for many tests can greatly reduce statistical power. Filtering out potentially uninformative features before running statistical tests can help address this problem, although this can also have unexpected effects (Bourgon et al., 2010). Importantly, this filtering must be independent of the test statistic evaluated (referred to as Independent Filtering). For instance, hard cut-offs for the prevalence and abundance of taxa across samples, and not within one group compared with another, are commonly used to exclude rare taxa (Schloss, 2020). This data filtering could be especially important for microbiome datasets because they are often extremely sparse. Nonetheless, it remains unclear whether filtering rare taxa has much effect on DA results in practice.

Another contentious area is regarding which statistical distributions are most appropriate for analyzing microbiome data. Statistical frameworks based on a range of distributions have been developed for modelling read count data. For example, DESeq2 (Love et al., 2014) and edgeR (Robinson and Oshlack, 2010) are both tools that assume normalized read counts follow a negative binomial distribution. To identify differentially abundant taxa, a null and alternative hypothesis are compared for each taxon. The null hypothesis is that the same parameters for the negative binomial solution explain the distribution of taxa across all sample groupings. The alternative hypothesis is that different parameters are needed to account for differences between sample groupings. If the null hypothesis can be rejected for a specific taxon then it is considered differentially abundant. This idea is the foundation of distribution-based DA tests, including other methods such as corncob (Martin et al., 2020) and metagenomeSeq (Paulson et al., 2013), which model microbiome data with the beta-binomial and zero-inflated Gaussian distributions, respectively.

Compositional data analysis (CoDa) methods represent an alternative to these approaches. It has recently become more widely appreciated that sequencing data are compositional (Gloor et al., 2017) meaning that sequencing only provides information on the relative abundance of features and that each feature is dependent on the relative abundance of all other features. This characteristic means that false inferences are commonly made when standard methods, intended for absolute abundances, are used with taxa relative abundances. CoDa methods circumvent this issue by reframing the focus to ratios of taxa relative abundances (Aitchison, 1982; Morton et al., 2019). The difference between CoDa methods considered in this paper is what quantity is used as the denominator, or the reference, for the transformation. The centred log-ratio (CLR) transformation is a CoDa approach that uses the geometric mean of the relative abundance of all taxa within a sample as the reference for that sample. An extension of this approach is implemented in the tool ALDEx2 (Fernandes et al., 2014) . The additive log-ratio transformation is an alternative approach where the reference is the relative abundance of a single taxon, which should be present with low variance in read counts across samples. ANCOM is one tool that implements this additive log-ratio approach (Mandal et al., 2015).

Evaluating the numerous options for analyzing microbiome data outlined above has proven difficult. This is largely because there are no gold standards to compare with DA tool results. Simulating datasets with specific taxa that are differentially abundant is a partial solution to this problem, but it is imperfect. For example, it has been noted that parametric simulations can result in circular arguments for specific tools (Hawinkel et al., 2019). It is unsurprising that distribution-based methods perform best when applied to simulated data based on that distribution. Nonetheless, simulated data with no expected differences has been valuable for evaluating the false discovery rate (FDR) of these methods. Based on this approach it has become clear that many of the methods output unacceptably high numbers of false positives (Calgaro et al., 2020; Feijen et al., 2016; Thorsen et al., 2016; Weiss et al., 2017). Similarly, based on simulated datasets with spiked taxa it has been shown that these methods can drastically vary in statistical power as well (Hawinkel et al., 2019; Thorsen et al., 2016).

Although these general observations have been well substantiated, there is less agreement regarding the performance of tools across evaluation studies. Certain observations have been reproducible, such as the higher FDR of edgeR and metagenomeSeq. Similarly, ALDEx2 has been repeatedly shown to have low power to detect differences (Calgaro et al., 2020; Hawinkel et al., 2019). In contrast, both ANCOM and limma voom (Law et al., 2014; Ritchie et al., 2015) have been implicated as both accurately and poorly controlling the FDR, depending on the study (Calgaro et al., 2020; Hawinkel et al., 2019; Weiss et al., 2017). To further complicate comparisons, different sets of tools and dataset types have been analyzed across evaluation studies. This means that, on some occasions, the best performing method in one evaluation is missing from another. In addition, certain popular microbiome-specific methods, such as MaAsLin2 (Mallick et al., 2021), have been missing from past evaluations. Finally, many evaluations limit their analysis to a small number of datasets that do not represent the breadth of datasets found in 16S rRNA gene sequencing studies.

Given the inconsistencies across these studies it is important that additional, independent evaluations be performed to elucidate the performance of current DA methods. Accordingly, herein we have conducted additional evaluations of common DA tools across 38 two-group 16S rRNA gene datasets. We first present the concordance of the methods on these datasets to investigate how consistently the methods cluster and perform in general, with and without the removal of rare taxa. Next, based on artificially subsampling the datasets into two groups where no differences are expected, we present the observed FDR for each DA tool. Lastly, we present an evaluation of how consistent biological interpretations would be across diarrheal datasets depending on which tool was applied. Our work enables improved assessment of these DA tools and highlights which key recommendations made by previous studies hold in an independent evaluation. Furthermore, our analysis shows various characteristics of DA tools that authors can use to evaluate published literature within the field.

## Methods

### Code and data availability

All code used for processing and analyzing the data is available in this GitHub repository: https://github.com/nearinj/Comparison_of_DA_microbiome_methods. The processed datasets and metadata files analyzed in this study are available on figshare: https://figshare.com/articles/dataset/16S_rRNA_Microbiome_Datasets/14531724. The accessions and/or locations of the raw data for each tested dataset are listed in **Supplementary Table 1**.

**Table 1:** Differential abundance tools compared in this study.

| Tool | Input | Norm. | Trans. | Distribution | Covariates | Random Effects | Hypothesis test | FDR Corr. | CoDa | Dev. For |
| --- | --- | --- | --- | --- | --- | --- | --- | --- | --- | --- |
| ALDEx2 | Counts | None | CLR | Dirichlet-multinomial | Yes* | No | Wilcoxon rank-sum | Yes | Yes | RNA-seq, 16S, MGS |
| ANCOM-II | Counts | None | ALR | Non-parametric | Yes | Yes | Wilcoxon rank-sum | Yes | Yes | MGS |
| corncob | Counts | None | None | Beta-binomial | Yes | No | Wald (default) | Yes | No | 16S, MGS |
| DESeq2 | Counts | Modified RLE (default is RLE) | None | Negative binomial | Yes | No | Wald (default) | Yes | No | RNA-seq, 16S, MGS |
| edgeR | Counts | RLE (default is TMM) | None | Negative binomial | Yes* | No | Exact | Yes | No | RNA-seq |
| LEFse | Rarefied relative abundance | TSS | None | Non-parametric | Subclass factor only | No | Kruskal-Wallis | No | No | 16S, MGS |
| MaAsLin2 | Counts | TSS | AST (default is log) | Normal (default) | Yes | Yes | Wald | Yes | No | MGS |
| MaAsLin2 (rare) | Rarefied counts | TSS | AST (default is log) | Normal (default) | Yes | Yes | Wald | Yes | No | MGS |
| metagenomeSeq | Counts | CSS | Log | Zero-inflated log-Normal | Yes | No | Moderated t | Yes | No | 16S, MGS |
| limma voom (TMM) | Counts | TMM | Log; Precision weighting | Normal (default) | Yes | Yes | Moderated t | Yes | No | RNA-seq |
| limma voom (TMMwsp) | Counts | TMMwsp | Log; Precision weighting | Normal (default) | Yes | Yes | Moderated t | Yes | No | RNA-seq |
| t-test (rare) | Rarefied Counts | None | None | Normal | No | No | Welch's t-test | Yes | No | N/A |
| Wilcoxon (CLR) | CLR abundances | None | CLR | Non-parametric | No | No | Wilcoxon rank-sum | Yes | Yes | N/A |
| Wilcoxon (rare) | Rarefied counts | None | None | Non-parametric | No | No | Wilcoxon rank-sum | Yes | No | N/A |

\* The tool supports additional covariates if they are provided. ANCOM-II automatically performs ANOVA in this case, ALDEx2 requires that users select the test, and edgeR requires use of a different function (glmFit or glmQLFit instead of exactTest). Abbreviations: ALR, additive log-ratio; AST, arcsine square-root transformation; CLR, centered log-ratio; CoDa, compositional data analysis; CSS, cumulative sum scaling; FDR Corr., false-discovery rate correction; MGS, metagenomic sequencing; RLE, relative log expression; TMM, trimmed mean of M-values; Trans., transformation; TSS, total sum scaling

### Dataset processing

Thirty-eight different datasets were included in our main analyses for assessing the characteristics of microbiome differential abundance tools. Two additional datasets were also included for a comparison of differential abundance consistency across diarrhea-related microbiome datasets. All datasets presented herein have been previously published or are publicly available (Alkanani et al., 2015; Baxter et al., 2016; Chase et al., 2016; De Tender et al., 2015; Dinh et al., 2015; Douglas et al., 2018; Dranse et al., 2018; Duvallet et al., 2017; Frère et al., 2018; Gonzalez et al., 2018; Goodrich et al., 2014; Hoellein et al., 2017; Ji et al., 2015; Kesy et al., 2019; Lamoureux et al., 2017; Lozupone et al., 2013; McCormick et al., 2016; Mejía-León et al., 2014; Nearing et al., 2019; Noguera-Julian et al., 2016; Oberbeckmann et al., 2016; Oliveira et al., 2018; Papa et al., 2012; Pop et al., 2014; Rosato et al., 2020; Ross et al., 2015; Scheperjans et al., 2015; Scher et al., 2013; Schneider et al., 2017; Schubert et al., 2014; Singh et al., 2015; Son et al., 2015; Turnbaugh et al., 2009; Vincent et al., 2013; Wu et al., 2019; Yurgel et al., 2017; Zeller et al., 2014; Zhu et al., 2013) (**Supp. Table 1**). Most datasets were already available in table format with ASV or operational taxonomic unit abundances while a minority needed to be processed from raw sequences. These raw sequences were processed with QIIME 2 version 2019.7 (Bolyen et al., 2018) based on the Microbiome Helper standard operating procedure (Comeau et al., 2017). Primers were removed using cutadapt (Martin, 2011) and stitched together using the QIIME 2 VSEARCH (Rognes et al., 2016) *join-pairs* plugin. Stitched reads were then quality filtered using the *quality-filter* plugin and reads were denoised using Deblur (Amir et al., 2017) to produce amplicon sequence variants (ASVs). Abundance tables of ASVs for each sample were then output into tab-delimited files. Rarefied tables were also produced for each dataset, where the rarefied read depth was taken to be the lowest read depth of any sample in the dataset over 2000 reads (with samples below this threshold discarded).

### Differential abundance testing

We created a custom shell script (run_all_tools.sh) that ran each differential abundance tool on each dataset within this study. As input the script took a tab-delimited ASV abundance table, a rarefied version of that same table, and a metadata file that contained a column that split the samples into two groups for testing. This script also accepted a prevalence cut-off filter to remove ASVs below a minimum cut-off, which was set to 10% (i.e., ASVs found in fewer than 10% of samples were removed) for the filtered data analyses we present. Note that in a minority of cases a genus abundance table was input instead, in which case all options were kept the same. When the prevalence filter option was set, the script also generated new filtered rarefied tables based on an input rarefaction depth.

Following these steps, each individual differential abundance method was run on the input data using either the rarefied or non-rarefied table, depending on which is recommended for that tool. The workflow used to run each differential abundance tool (with run_all_tools.sh) is described below. The first step in each of these workflows was to read the dataset tables into R (version 3.6.3) with a custom script and then ensure that samples within the metadata and feature abundance tables were in the same order. An alpha-value of 0.05 was chosen as our significance cutoff and FDR-corrected p-values were used for methods that output p-values.

#### ALDEx2

We passed the non-rarefied feature table and the corresponding sample metadata to the *aldex* function from the ALDEx2 R package (Fernandes et al., 2014) which generated Monte Carlo samples of Dirichlet distribution for each sample, using a uniform prior, performed CLR transformation of each realization, and performed Wilcoxon tests on the transformed realizations. The function then returned the expected Benjamini-Hochberg (BH) FDR-corrected P-value was then returned for each feature based on the results across Monte Carlo samples.

#### ANCOM-II

We ran the non-rarefied feature table through the R ANCOM-II (Kaul et al., 2017; Mandal et al., 2015) (https://github.com/FrederickHuangLin/ANCOM) function *feature_table_pre_process*, which first examined the abundance table to identify outlier zeros and structural zeros (Kaul et al., 2017). Outlier zeros, identified by finding outliers in the distribution of taxon counts within each sample grouping, were ignored during differential abundance analysis and replaced with NA. Structural zeros, taxa that were absent in one grouping but present in the other, were ignored during data analysis and automatically called as differentially abundant. A pseudo count of 1 was then applied across the dataset to allow for log transformation. Using the main function *ANCOM*, all additive log-ratios for each taxon were then tested for significance using Wilcoxon rank-sum tests, and p-values were FDR-corrected using the BH method. ANCOM then applied a detection threshold as described in the original paper (Mandal et al., 2015), whereby a taxon was called as DA if the number of corrected p-values reaching nominal significance for that taxon was greater than 90% of the maximum possible number of significant comparisons.

#### corncob

We converted the metadata and non-rarefied feature tables into a phyloseq object, which we input to corncob’s *differentialTest* function (Martin et al., 2020). This function first converted the data into relative abundances and then fit each taxon abundance to a beta-binomial model, using logit link functions for both the mean and overdispersion. Because corncob models each of these simultaneously and performs both differential abundance and differential variability testing (Martin et al., 2020), we set the null overdispersion model to be the same as the non-null model so that only taxa having differential abundances were identified. Finally, the function performed significance testing, for which we chose Wald tests (with the default non-bootstrap setting), and we obtained BH FDR-corrected p-values as output.

#### DESeq2

We first passed the non-rarefied feature tables to the *DESeq* function (Love et al., 2014) with default settings, except that instead of the default relative log expression (also known as the median-of-ratios method) the estimation of size factors was set to use “poscounts”, which calculates a modified relative log expression that helps account for features missing in at least one sample. The function performed three steps: (1) estimation of size factors, which are used to normalize library sizes in a model-based fashion; (2) estimation of dispersions from the negative binomial likelihood for each feature, and subsequent shrinkage of each dispersion estimate towards the parametric (default) trendline by empirical Bayes; (3) fitting each feature to the specified class groupings with negative binomial generalized linear models and performing hypothesis testing, for which we chose the default Wald test. Finally, using the *results* function, we obtained the resulting BH FDR-corrected p-values.

#### edgeR

Using the phyloseq_to_edgeR function (https://joey711.github.io/phyloseq-extensions/edgeR.html), we added a pseudocount of 1 to the non-rarefied feature table and used the function *calcNormFactors* from the edgeR R package (Robinson and Oshlack, 2010) to compute relative log expression normalization factors. Negative binomial dispersion parameters were then estimated using the functions *estimateCommonDisp* followed by *estimateTagwiseDisp* to shrink feature-wise dispersion estimates through an empirical Bayes approach. We then used the *exactTest* for negative binomial data (Robinson and Oshlack, 2010) to identify features that differ between the specified groups. The resulting p-values were then corrected for multiple testing with the BH method with the function *topTags*.

#### LEfSe

The rarefied feature table was first converted into LEfSe format using the LEfSe script *format_input*.*py* (Segata et al., 2011). We then ran LEfSe on the formatted table using the *run_lefse*.*py* script with default settings and no subclass specifications. Briefly, this command first normalized the data using total sum scaling, which divides each feature count by the total library size. Then it performed a Kruskal-Wallis (which in our two-group case reduces to the Wilcoxon rank-sum) hypothesis test to identify potential differentially abundant features, followed by linear discriminant analysis of class labels on abundances to estimate the effect sizes for significant features. From these, only those features with scaled logarithmic linear discriminant analysis scores above the threshold score of 2.0 (default) were called as differentially abundant.

#### limma voom

We first normalized the non-rarefied feature table using the edgeR *calcNormFactors* function, with either the trimmed mean of M-values (TMM) or TMM with singleton pairing (TMMwsp) option. During this step (for both options), a single sample was chosen to be a reference sample using upper-quartile normalization. This step failed in some highly sparse abundance tables, in which cases, we instead chose the sample with the largest sum of square-root transformed feature abundances to be the reference sample. After normalization, we used the limma R package (Ritchie et al., 2015) function *voom* to convert normalized counts to log_2_-counts-per-million and assign precision weights to each observation based on the mean-variance trend. We then used the functions *lmFit, eBayes*, and *topTable* to fit weighted linear regression models, perform tests based on an empirical Bayes moderated t-statistic (Phipson et al., 2016) and obtain BH FDR-corrected p-values.

#### MaAsLin2

We entered either a rarefied or non-rarefied feature table into the main *Maaslin2* function within the Maaslin2 R package(Mallick et al., 2021). We specified arcsine square-root transformation as in the package vignette (instead of the default log) and total sum scaling normalization. For consistency with other tools, we specified no random effects and turned off default standardization. The function fit a linear model to each feature’s transformed abundance on the specified sample grouping, tested significance using a Wald test, and output BH FDR-corrected p-values.

#### metagenomeSeq

We first entered the counts and sample information to the function *newMRexperiment* from the metagenomeSeq R package (Paulson et al., 2013). Next, we used *cumNormStat* and *cumNorm* to apply cumulative sum-scaling normalization, which attempts to normalize sequence counts based on the lower-quartile abundance of features. We then used *fitFeatureModel* to fit normalized feature counts with zero-inflated log-normal models (with pseudo-counts of 1 added prior to log_2_ transformation) and perform empirical Bayes moderated t-tests, and *MRfulltable* to obtain BH FDR-corrected p-values.

#### t-test

We applied total sum scaling normalization to the rarefied feature table, and performed an unpaired Welch’s t-test for each feature to compare the specified groups. We corrected the resulting p-values for multiple testing with the BH method.

#### Wilcoxon test

Using raw feature abundances in the rarefied case, and CLR-transformed abundances (after applying a pseudocount of 1) in the non-rarefied case, we performed Wilcoxon rank-sum tests for each feature to compare the specified sample groupings. We corrected the resulting p-values with the BH method.

### Comparing numbers of significant hits between tools

We compared the number of significant ASVs each tool identified in 38 different datasets. Each tool was run as described above using default settings with some modifications suggested by the tool authors, as noted above. A heatmap representing the number of significant hits found by each tool was constructed using the pheatmap R package (Kolde, 2012). Spearman correlations between the number of significant ASVs identified by a tool and the following dataset characteristics were computed using the cor.test function in R: sample size, Aitchison’s distance effect size as computed using a PERMANOVA test (adonis; vegan) (Dixon, 2003), sparsity, mean sample ASV richness, median sample read depth, read depth range between samples and the coefficient of variation for read depth within a dataset.

### Cross-tool, within-study differential abundance consistency analysis

We compared the consistency between different tools within all datasets by pooling all ASVs identified as being significant by at least one tool in the 38 different datasets. The number of methods that identified each ASV as differentially abundant were then tallied.

### False positive analysis

To estimate the false positives a method might produce during data analysis, eight datasets were selected for analysis. These datasets were chosen based on having the largest sample sizes, while also being from diverse environment types. In each dataset, the most frequent sample group was chosen for analysis to help ensure similar composition among samples tested. Within this grouping, random labels of either case or control were assigned to samples and the various differential abundance methods were tested on them. This was repeated 10 times for each dataset and each tool tested with an additional 90 replications for all tools except for ALDEx2, ANCOM-II and corncob due to high resource requirements. After analysis was completed, the number of differentially abundant ASVs identified by each tool was assessed at an alpha value of 0.05.

### Cross-study differential abundance consistency analysis

Two additional datasets were acquired to bring the number of diarrhea-related datasets to five. The ASVs in each of these datasets were previously taxonomically classified and so we used these classifications to collapse all feature abundances to the genus level. Note that taxonomic classification was performed using several different methods, which represents another source of technical variation. We excluded unclassified and *sensu stricto*-labelled genus levels. We then ran all differential abundance tools on these datasets at the genus level. These comparisons were between the diarrhea and non-diarrhea sample groups. The same processing workflow was used for the supplementary obesity dataset comparison as well.

For each tool and study combination, we determined which genera were significantly different at an alpha of 0.05 (where relevant). For each tool we then tallied up the number of times each genus was significant, i.e., how many datasets each genus was significant in based on a given tool. The null expectation distributions of these counts per tool were generated by randomly sampling genera from each dataset for 100,000 replicates. The probability of sampling a genus (i.e., calling it significant) was set to be equal to the proportion of actual significant genera. For each replicate we tallied up the number of times each genus was sampled across datasets. We then compared the observed and expected distributions of the number of studies each genus was found to be significant in. Note that to simplify this analysis we ignored the directionality of the significance (e.g., whether it was higher in case or control samples). We excluded genera never found to be significant. We performed bootstrap Kolmogorov-Smirnov tests (10,000 replicates) using the *ks*.*boot* function from the Matching R package (Sekhon, 2011) to compare the expected and observed distributions for each tool.

## Results

### Microbiome differential abundance methods produce a highly variable number of significant ASVs within the same microbiome datasets

To investigate how different DA tools impact biological interpretations across microbiome datasets, we tested 14 different differential abundance testing approaches (**Table 1**) on 38 different microbiome datasets. These datasets corresponded to a range of environments, including the human gut, plastisphere, freshwater lakes, and urban environments (**Supp. Table 1**). The features in these datasets corresponded to both ASVs and clustered operational taxonomic units, but we refer to them all as ASVs below for simplicity.

We also investigated how prevalence filtering each dataset prior to analysis impacted the observed results. We chose to either use no prevalence filtering (**Fig. 1A**) or a 10% prevalence filter that removed any ASVs found in fewer than 10% of samples within each dataset (**Fig. 1B**).

**Figure 1:**
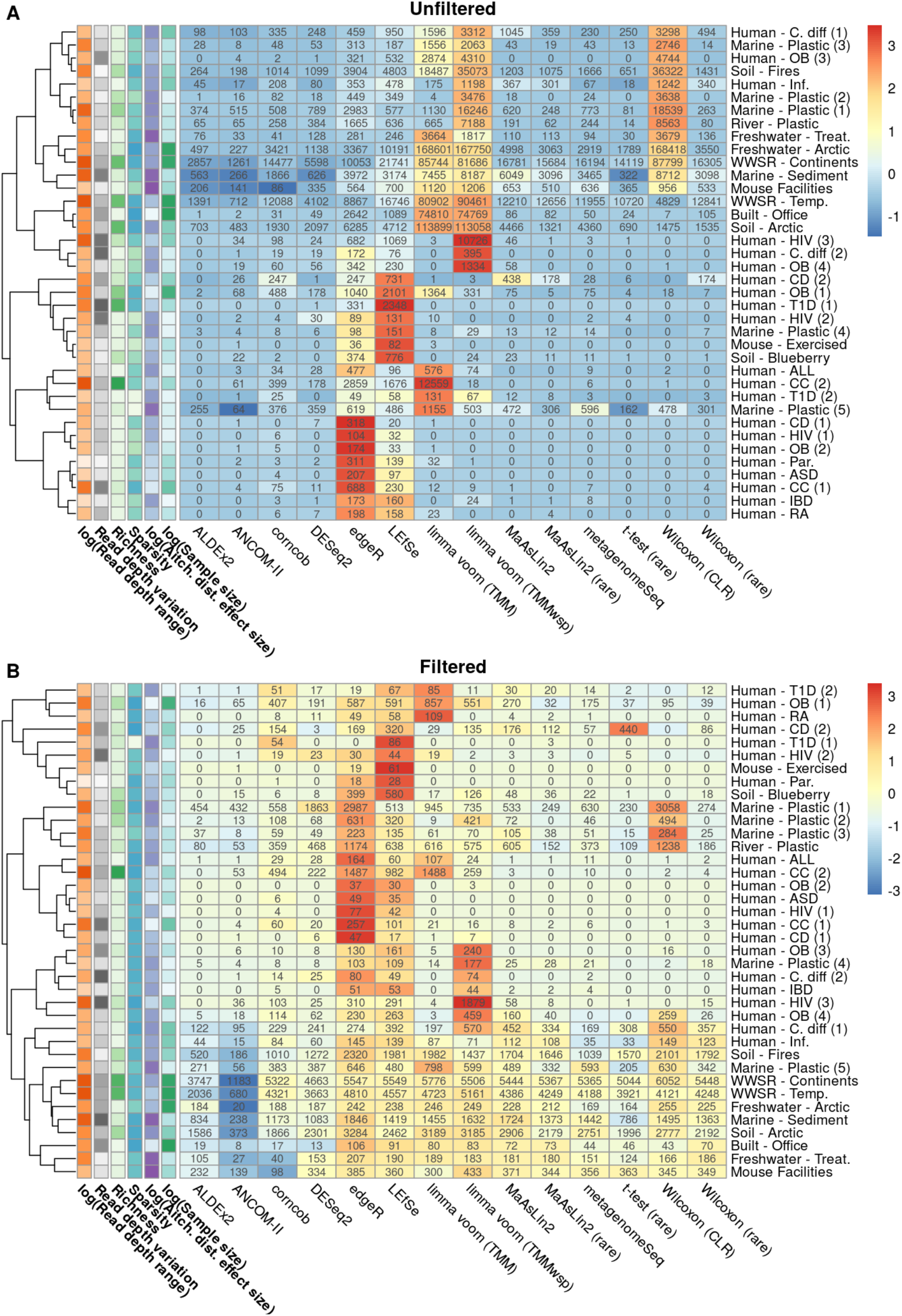
Variation in the proportion of significant features depending on the differential abundance method and dataset. Heatmaps indicate the numbers of significant amplicon sequence variants (ASVs) identified in each dataset by the corresponding tool based on (A) unfiltered data and (B) 10% prevalence-filtered data. Cells are coloured based on the standardized (scaled and mean centred) percentage of significant ASVs for each dataset. Additional coloured cells in the left-most six columns indicate the dataset characteristics we hypothesized could be driving variation in these results. Darker colours indicate higher values in these six columns. Datasets were hierarchically clustered based on Euclidean distances using the complete method.

We found that in both the filtered and unfiltered analyses the percentage of significant ASVs identified by each DA method varied widely across datasets, with means ranging from 3.8-32.5% and 0.8-40.5%, respectively. Interestingly, we found that many tools behaved differently between datasets. Specifically, some tools identified the most features in one dataset while identifying only an intermediate number in other datasets. This was especially evident in the unfiltered datasets (**Fig. 1A**).

To investigate possible factors driving this variation we examined how the number of ASVs identified by each tool correlated with several variables. These variables included dataset richness, variation in sequencing depth between samples, dataset sparsity, and Aitchison’s distance effect size (based on PERMANOVA tests). As expected, we found that all tools positively correlated with the effect size between test groups with rho values ranging between 0.35-0.72 with unfiltered data (**Fig. 2A**) and 0.31-0.52 for filtered data (**Fig. 2B**). We also found in the filtered datasets that the number of features found by all tools significantly correlated with the median read depth, range in read depth, and sample size. There was much less consistency in these correlations across the unfiltered data. For instance, only the t-test, both Wilcoxon methods, and both limma voom methods correlated significantly with the range in read depth **(Fig. 2B**). We also found that edgeR was negatively correlated with mean sample richness in the unfiltered analysis.

**Figure 2:**
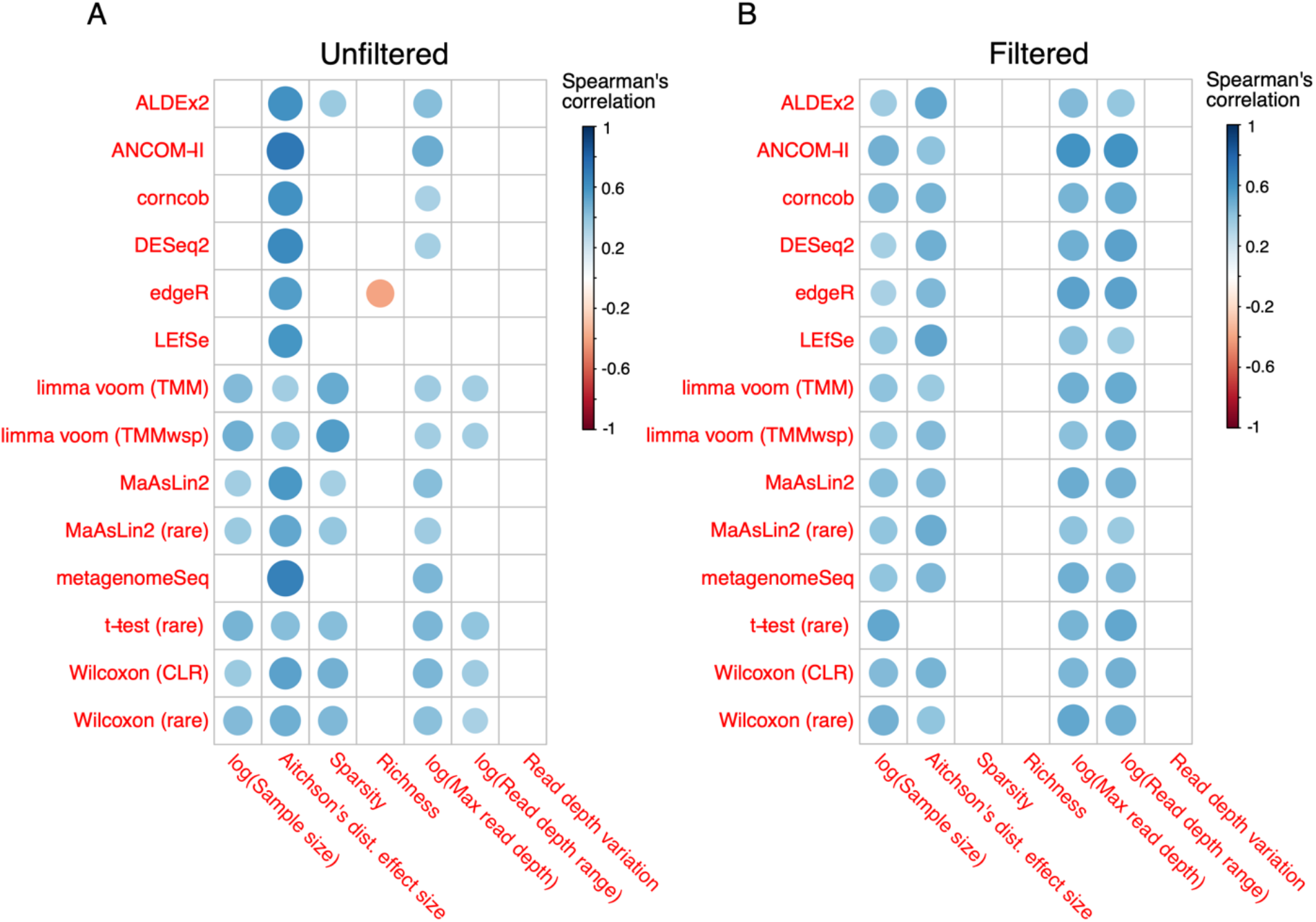
Dataset characteristics associated with percentage of significant amplicon sequence variants. The correlation coefficients (Spearman’s rho) are displayed by size and color. These correspond to the dataset characteristics correlated with the percentage of significant amplicon sequence variants identified by that tool per dataset. Only significant correlations (p < 0.05) are displayed.

Despite the variability of tool performance between datasets, we did find that several tools tended to identify more significant hits (**Supp. Fig 1C-D**). In the unfiltered datasets, we found that limma voom (TMMwsp; mean: 40.5% / TMM; mean: 29.7%), Wilcoxon (CLR; mean: 30.7%), LEfSe (mean: 12.6%), and edgeR (mean: 12.4%) tended to find the largest number of significant ASVs compared with other methods. Interestingly, in a few datasets, such as the Human-ASD and Human-OB (2) datasets, edgeR found a significantly higher proportion of significant ASVs than any other tool. In addition, we found that limma voom (TMMwsp) found the majority of ASVs to be significant (73.5%) in the Human-HIV (3) dataset while the other tools found 0-11% ASVs to be significant (**Fig. 1A**). Similarly, we found that both limma voom methods identified over 99% of ASVs to be significant in several cases such as the Built-Office and Freshwater-Arctic datasets. We found similar, although not as extreme, trends with LEfSe where in some datasets, such as the Human-T1D (1) dataset, the tool found a much higher percentage of significant hits (3.5%) compared with all other tools (0-0.4%). This observation is most likely a result of LEfSe filtering significant features by effect size rather than using FDR correction to reduce the number of false positives. We found that two of the three compositionally aware methods we tested identified fewer significant ASVs than the other tools tested. Specifically, ALDEx2 (mean: 1.4%) and ANCOM-II (mean: 0.8%) identified the fewest significant ASVs. We found the conservative behavior of these tools to be consistent across all 38 datasets we tested.

Overall, the results based on the filtered tables were similar, although there was a smaller range in the number of significant features identified by each tool. All tools except for ALDEx2 found a lower number of total significant features when compared with the unfiltered dataset (**Supp. Fig 1C-D**). As with the unfiltered data, ANCOM-II was the most stringent method (mean: 3.8%), while edgeR (mean: 32.5%), LEfSe (mean: 27.6%), limma voom (TMMwsp; mean: 27.3% / TMM; mean: 23.5%), and Wilcoxon (CLR; mean: 25.4%) tended to output the highest numbers of significant ASVs (**Fig. 1B**).

Finally, we examined the mean relative abundance of the features identified by each tool to determine whether tools may be biased toward the identification of highly abundant features. We found that both ALDEx2 (median: 0.013%), ANCOM-II (median: 0.024%) and to a lesser degree DESeq2 (median: 0.006%) tended to find significant features that were higher in relative abundance in the unfiltered datasets. A similar trend for ALDEx2 (median: 0.011%) and ANCOM-II (median: 0.029%) was also found in the filtered datasets (**Supp. Fig 1A-B**).

### High variability of overlapping significant ASVs

We next investigated the overlap in significant ASVs across tools within each dataset. Based on the unfiltered data, we found that both limma voom methods identified similar sets of significant ASVs that were different from those of most other tools (**Fig. 3A**). However, we also found that many of the ASVs identified by the limma voom methods were also identified as significant based on the Wilcoxon (CLR) approach, despite these being highly methodologically distinct tools. Furthermore, the two Wilcoxon test approaches had different consistency profiles despite using the same hypothesis test. In contrast, we found that both MaAsLin2 approaches had similar consistency profiles, although the non-rarefied method found slightly lower-ranked features. We also found that the most conservative tools, ALDEx2 and ANCOM-II, primarily identified features that were also identified by almost all other methods. In contrast, edgeR and LEfSe, two tools that often identified the most significant ASVs, output the highest percentage of ASVs that were not identified by any other tool: 12.1% and 11.1%, respectively. Corncob, metagenomeSeq, and DESeq2 identified ASVs at more intermediate consistency profiles.

**Figure 3:**
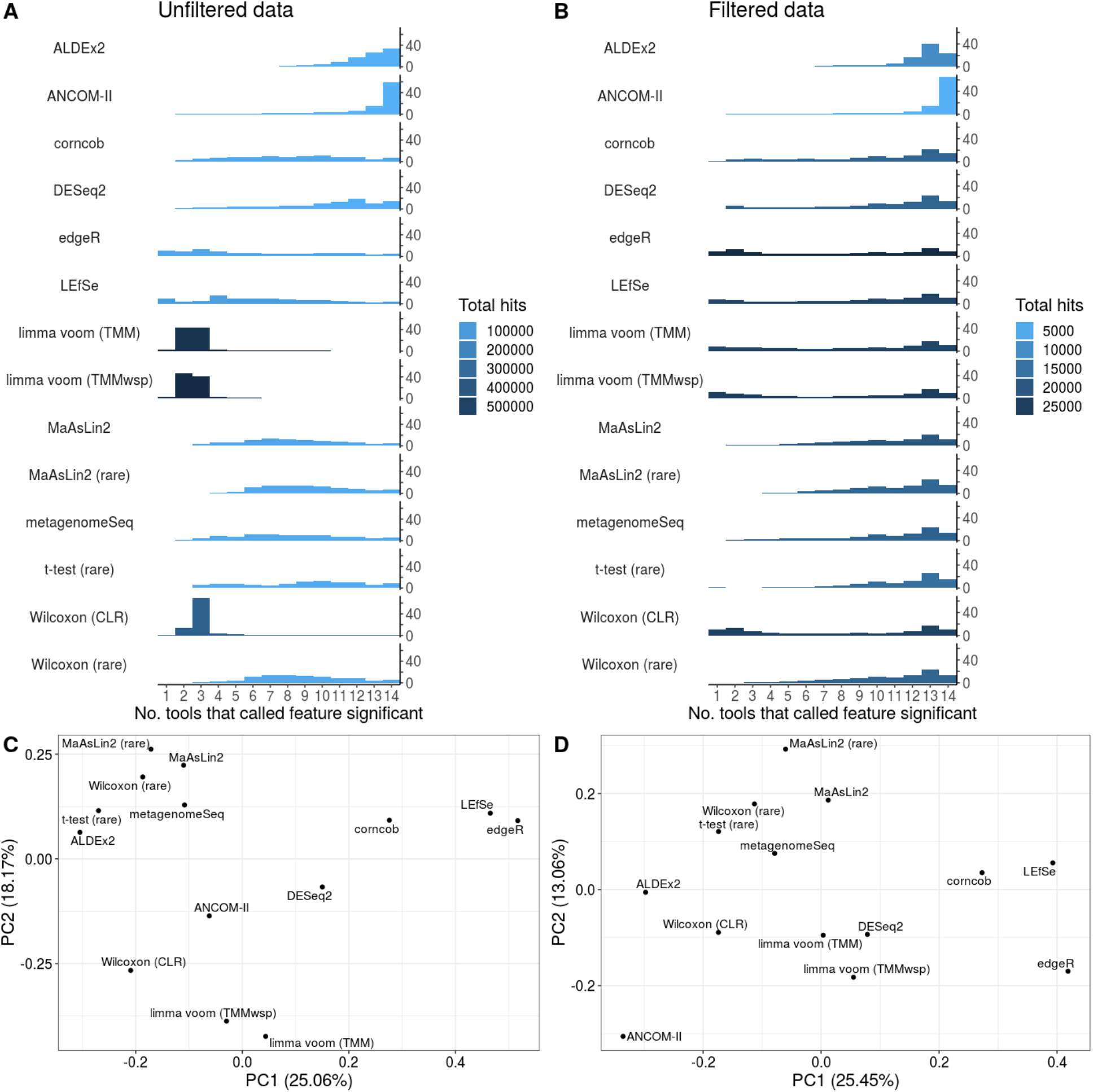
Overlap of significant features across tools and tool clustering. (A and B) The number of tools that called each feature significant, stratified by features called by each individual tool for the (A) unfiltered and (B) 10% prevalence-filtered data. The features correspond to the amplicon sequence variants (and operational taxonomic units) from all 38 tested datasets. Results are shown as a percentage of all ASVs identified by each tool. The total number of significant features identified by each tool is indicated by the bar colors. (C and D) Plots are displayed for the first two principal coordinates (PCs) for both (C) non-prevalence-filtered and (D) 10% prevalence-filtered data. These plots are based on the mean inter-tool Jaccard distance across the 38 main datasets that we analyzed, computed by averaging over the inter-tool distance matrices for all individual datasets in order to weight each dataset equally.

The overlap in significant ASVs based on the prevalence-filtered data was similar overall to the unfiltered data results (**Fig. 3B**). One important exception was that the limma voom approaches identified a much higher proportion of ASVs that were also identified by most other tools, compared with the unfiltered data. Nonetheless, similar to the unfiltered data results, the Wilcoxon (CLR) significant ASVs displayed a bimodal distribution and a strong overlap with limma voom methods. We also found that overall, the proportion of ASVs consistently identified as significant by more than 12 tools was much higher in the filtered data (mean: 38.5; SD: 15.8) compared with the unfiltered data (mean: 17.3; SD: 22.1). In contrast with the unfiltered results, corncob, metagenomeSeq, and DESeq2 had lower proportions of ASVs at intermediate consistency ranks. However, ALDEx2 and ANCOM-II once again produced significant ASVs that largely overlapped with other tools.

The above analyses summarized the variation in tool performance across datasets, but it is difficult to discern which tools performed most similarly from these results alone. To identify overall similarly performing tools we conducted principal coordinates analysis based on the Jaccard distance between significant sets of ASVs (**Fig. 3C, 3D**). One clear trend for both unfiltered and filtered data is that edgeR and LEfSe cluster together and are separated from other methods on the first principal coordinate. Interestingly, corncob, which is a methodologically distinct approach, also clusters relatively close to these two methods on the first PC. The major outliers on the second principal coordinate differ depending on whether the data was prevalence-filtered. For the unfiltered data, the main outliers are the limma voom methods, followed by Wilcoxon (CLR; **Fig. 3C**). In contrast, ANCOM-II is the sole major outlier on the second principal component based on filtered data (**Fig. 3D**). These visualizations highlight the major tool clusters based on the resulting sets of significant ASVs. However, the percentage of variation explained by the top two components is relatively low in each case, which means that substantial information regarding potential tool clustering is missing from these panels (**Supp**.

**Fig. 2** and **Supp. Fig. 3**). For instance, ANCOM-II and corncob are major outliers on the third and fourth principal coordinates, respectively, of the unfiltered data analysis, which highlights the uniqueness of these methods.

### False discovery rate of microbiome differential abundance tools depends on the dataset

We next investigated the FDR of each DA tool across eight datasets. For each dataset we selected the most frequently sampled group and randomly reassigned them as case or control samples. Each DA tool was then run on those samples and results were compared. This was repeated a total of 10 times for each filtered dataset and 100 times for each unfiltered dataset (except for corncob, ANCOM-II and ALDEx2). We used a higher number of replicates for the unfiltered datasets because they were less stable across replicates. The percentage of significant ASVs after BH multiple-test correction was relatively low for most tested tools on the filtered data (**Fig. 4B**). Two outliers were edgeR (mean: 10.3%; SD: 9.0%) and LEfSe (mean: 4.4%; SD: 1.2%), which consistently identified more significant hits compared with other tools (range of other tool means: 0% -2.2%). Both limma voom methods output highly variable percentages of significant ASVs, especially based on the unfiltered data (**Fig. 4A**). In particular, in 5/8 of the unfiltered datasets, the limma voom methods identified more than 5% of ASVs as significant on average. Interestingly, while these two methods exhibited similar performance overall, the performance within the unfiltered Freshwater-Treatment dataset was highly different between the methods with the TMMwsp method identifying 0.001% and the TMM method identifying 9.0%. Only ALDEx2 and the t-test (rare) approach consistently identified no ASVs as significantly different in this analysis.

**Figure 4:**
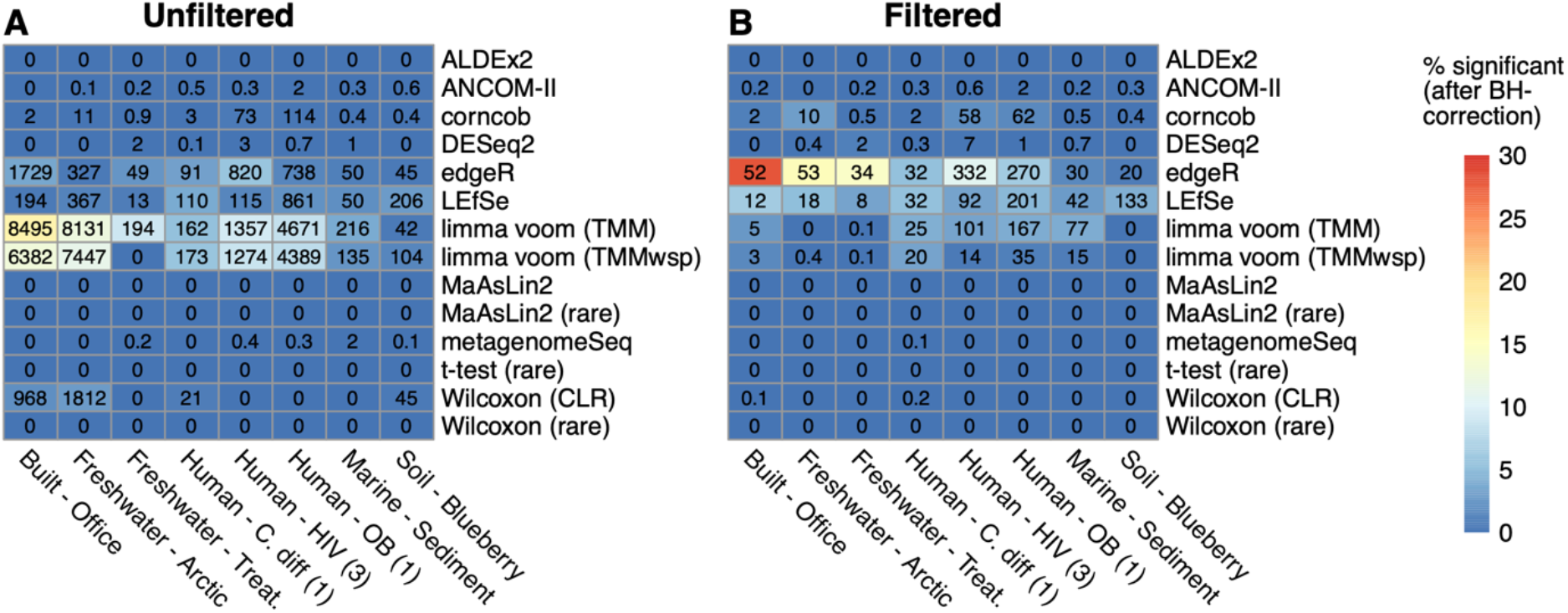
Variation in false discovery rate across the tested differential abundance tools in the context of simple simulations. Heatmap of the percentage and number of significant amplicon sequence variants (ASVs) identified by each tool in eight simulation datasets based on applying (A) no prevalence filter and (B) a 10% prevalence filter. Cell colours indicate the percentage and the number of significant ASVs is written in each cell. Mean numbers higher than one were rounded to the nearest integer for visualization. Significant ASVs were identified after applying the Benjamini-Hochberg (BH) false discovery rate procedure and then using a cut-off of 0.05.

Overall, we found that the raw numbers of significant ASVs were lower in the filtered dataset than in the unfiltered data (as expected due to many ASVs being filtered out), and that most tools identified only a small percentage of significant ASVs, regardless of filtering procedure. The exceptions were the two limma voom methods, which had high FDRs with unfiltered data, and edgeR and LEfSe, which had high FDRs on the filtered data. Although these tools stand out on average, we also observed that in several replicates on the unfiltered datasets, the Wilcoxon (CLR) approach identified almost all features as statistically significant (**Supp. Fig. 4)**. This was also true for both limma voom methods, which highlights that a minority of replicates are driving up the average FDR of these methods. Such extreme values were not observed for the filtered data (**Supp. Fig 5**).

Investigation into these outlier replicates for the Wilcoxon (CLR) approach revealed that the mean differences in read depth between the two tested groups were consistently higher in replicates in which 30% or more of ASVs were significant. Interestingly, this pattern was absent when examining replicates for the limma voom methods (**Supp. Fig 6**).

### Tools vary in how consistently they identify the same significant genera within diarrhea case-control datasets

Separate from the above analysis comparing consistency between tools on the same dataset, we next investigated whether certain tools provide more consistent signals across datasets of the same disease. This analysis focused on the genus-level across tools to help limit inter-study variation. We specifically focused on diarrhea as a phenotype, which has been shown to exhibit a strong effect on the microbiome and to be relatively reproducible across studies (Duvallet et al., 2017).

We acquired five datasets for this analysis representing the microbiome of individuals with diarrhea compared with individuals without diarrhea (see Methods). We ran all DA tools on each individual filtered dataset. Similar to our ASV-level analyses, the tools substantially varied in terms of the number of significant genera identified. For instance, ALDEx2 identified a mean of 17.6 genera as significant in each dataset (SD: 17.4), while edgeR identified a mean of 46.0 significant genera (SD: 12.9). Tools that identify more genera as significant in general are accordingly more likely to identify genera as consistently significant compared with tools with fewer significant hits. Accordingly, inter-tool comparisons of the number of times each genus was identified as significant would not be informative.

Instead, we analyzed the observed distribution of the number of studies that each genus was identified as significant in compared with the expected distribution given random data. This approach enabled us to compare the tools based on how much more consistently each tool performed relative to its own random expectation. For instance, on average edgeR identified significant genera more consistently across studies compared with ALDEx2 (mean numbers of datasets that genera were found in across studies were 1.67 and 1.54 for edgeR and ALDEx2, respectively). However, this observation was simply driven by the increased number of significant genera identified by edgeR. Indeed, when compared with the random expectation, ALDEx2 displayed a 1.35-fold increase (p < 0.0001) of consistency in calling significant genera in the observed data. In contrast, edgeR produced results that were only 1.10-fold more consistent compared with the random expectation (p = 0.02).

ALDEx2 and edgeR represent the extremes of how consistently tools identify the same genera as significant across studies, but there is a large range (**Fig. 5**). Notably, all tools were significantly more consistent than the random expectation across these datasets (p < 0.05) (**Table 2**). In addition to ALDEx2, the other top performing approaches based on this evaluation included both MaAsLin2 workflows, limma voom (TMM), and ANCOM-II.

**Figure 5:**
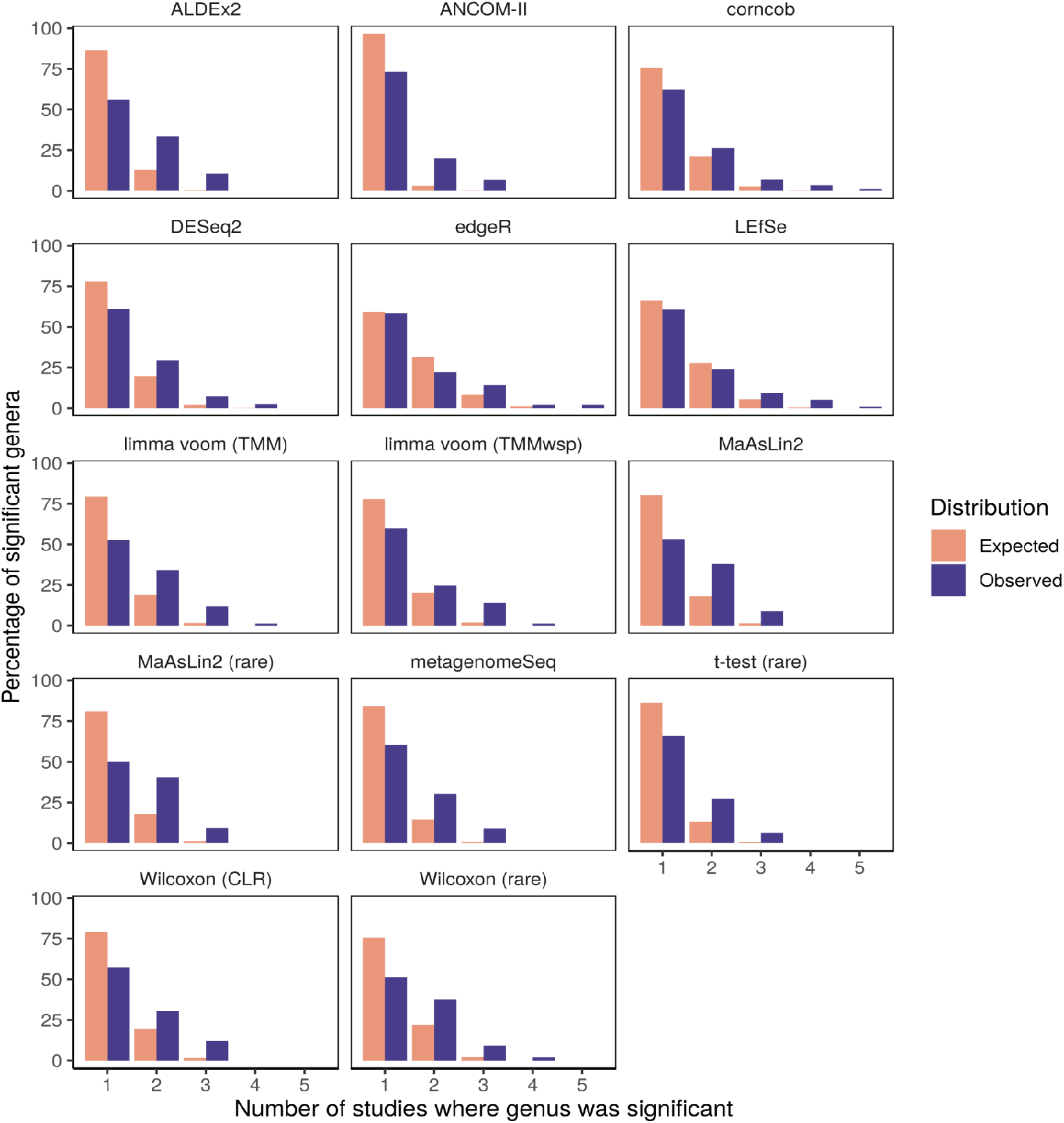
Observed consistency of significant genera across diarrhea datasets is higher than the random expectation overall. These barplots illustrate the distributions of the number of studies for which each genus was identified as significant (excluding genera never found to be significant). The random expectation distribution is based on replicates of randomly selecting genera as significant and then computing the consistency across studies.

**Table 2:** Results of Kolmogorov-Smirnov tests comparing observed and expected consistency in differentially abundant genera across five diarrhea datasets.

| Tool | No. sig. genera | Max overlap | Mean exp. | Mean obs. | Fold diff. | D | p |
| --- | --- | --- | --- | --- | --- | --- | --- |
| ALDEx2 | 57 | 3 | 1.14 | 1.544 | 1.354 | 0.304 | < 1e-4 |
| MaAsLin2 (rare) | 74 | 3 | 1.204 | 1.595 | 1.325 | 0.309 | < 1e-4 |
| limma voom (TMM) | 76 | 4 | 1.223 | 1.618 | 1.323 | 0.267 | < 1e-4 |
| ANCOM-II | 15 | 3 | 1.033 | 1.333 | 1.29 | 0.234 | 0.0009 |
| MaAsLin2 | 79 | 3 | 1.211 | 1.557 | 1.286 | 0.271 | < 1e-4 |
| Wilcoxon (rare) | 88 | 4 | 1.27 | 1.625 | 1.28 | 0.244 | < 1e-4 |
| metagenomeSeq | 66 | 3 | 1.164 | 1.485 | 1.276 | 0.239 | < 1e-4 |
| limma voom (TMMwsp) | 85 | 4 | 1.238 | 1.565 | 1.264 | 0.181 | < 1e-4 |
| Wilcoxon (CLR) | 82 | 3 | 1.225 | 1.549 | 1.264 | 0.218 | < 1e-4 |
| t-test (rare) | 62 | 3 | 1.145 | 1.403 | 1.225 | 0.201 | 0.0001 |
| corncob | 87 | 5 | 1.276 | 1.552 | 1.216 | 0.136 | 0.0030 |
| DESeq2 | 82 | 4 | 1.245 | 1.512 | 1.214 | 0.17 | 0.0001 |
| LEfSe | 117 | 5 | 1.402 | 1.615 | 1.152 | 0.095 | 0.0300 |
| edgeR | 138 | 5 | 1.511 | 1.667 | 1.103 | 0.096 | 0.0210 |

We conducted a similar investigation across five obesity 16S datasets, which was more challenging to interpret due to the lower consistency in general (**Supp. Table 2**). Specifically, most significant genera were called in only a single study and only MaAslin2 (both with non-rarefied and rarefied data) and the t-test (rare) approaches performed significantly better than expected by chance (p < 0.05). The MaAsLin2 (rare) approach produced by far the most consistent results based on these datasets (fold difference: 1.23; p = 0.006).

## Discussion

Herein we have compared the performance of commonly used DA tools, primarily on actual 16S datasets. While it might be argued that differences in tool outputs are expected given that they test different hypotheses, we believe this perspective ignores how these tools are used in practice. In particular, these tools are frequently used interchangeably in the microbiome literature.

Accordingly, an improved understanding of the variation in DA method performance is crucial to properly interpret microbiome studies. We have illustrated here that these tools can produce substantially different results, which highlights that many biological interpretations based on microbiome data analysis are likely not robust to DA tool choice. Our findings should serve as a cautionary tale for researchers conducting their own microbiome data analysis and reinforce the need to honestly report the findings of a representative set of different analysis options to ensure robust results are reported. Despite the high variation across DA tool results, we were able to characterize several consistent patterns produced by various tools that researchers should keep in mind when assessing both their own results and results from published work.

Two major groups of DA tools could be distinguished by how many significant ASVs they tended to identify. We found that limma voom, edgeR, Wilcoxon (CLR), and LEfSe output a high number of significant ASVs on average. In contrast, ALDEx2 and ANCOM-II tended to identify only a relatively small number of ASVs as significant. We hypothesize that these latter tools are more conservative and have higher precision, but with a concomitant probable loss in sensitivity. This hypothesis is related to our observation that significant ASVs identified by these two tools tended to also be identified by almost all other differential abundance methods, which we interpret to be ASVs that are more likely to be true positives.

Given that ASVs commonly identified as significant are likely more reliable, it is noteworthy that significant ASVs in the unfiltered data tended to be called by fewer tools. This was particularly true for both limma voom approaches and the Wilcoxon (CLR) approach.

Although it is possible that many of these significant ASVs are incorrectly missed by other tools, it is more likely that these tools are simply performing especially poorly on unfiltered data due to several reasons, such as data sparsity.

This issue with the limma voom approaches was also highlighted by high false positive rates on several unfiltered randomized datasets, which agrees with a past FDR assessment of this approach (Hawinkel et al., 2019). It is also important to acknowledge that our randomized approach for estimating FDR is not a perfect representation of real data; that is, real sample groupings will likely contain some systematic differences in microbial abundances—although the effect size may be very small—whereas our randomized datasets should have none. Accordingly, identifying only a few significant ASVs under this approach is not necessarily proof that a tool has a low FDR in practice. However, tools that identified many significant ASVs in the absence of distinguishing signals likely also have high FDR on real data.

Two additional particularly problematic tools based on this analysis were edgeR and LEfSe. The edgeR method has been previously found to exhibit a high FDR on several occasions (Hawinkel et al., 2019; Thorsen et al., 2016) Although metagenomeSeq also has been flagged as such (Thorsen et al., 2016), that was not the case in our analysis. This agrees with a recent report that metagenomeSeq (using the zero-inflated log-normal approach, as we did) appropriately controlled the FDR, but exhibited low power (Lin and Peddada, 2020). There have been mixed results previously regarding whether ANCOM appropriately controls the FDR (Hawinkel et al., 2019; Weiss et al., 2017), but the results from our limited analysis suggest that this method is conservative and controls the FDR while potentially missing true positives.

Related to this point, we found that ANCOM-II performed better than average at identifying the same genera as significantly DA across five diarrhea-related datasets despite only identifying a mean of four genera as significant per dataset. Nonetheless, the ANCOM-II results were less consistent than ALDEx2, both MaAsLin2 workflows, and limma voom (TMM). The tools that produced the least consistent results across datasets (relative to the random expectation) included the t-test (rare) approach, LEfSe, and edgeR. The random expectation in this case was quite simplistic; it was generated based on the assumption that all genera were equally likely to be significant by chance. This assumption must be invalid to some degree simply because some genera are more prevalent than others across samples. Accordingly, it is surprising that the tools produced only marginally more consistent results than expected.

Although this cross-data consistency analysis was informative, it was interesting to note that not all environments and datasets are appropriate for this comparison. Specifically, we found that the consistency of significant genera across five datasets comparing obese and control individuals was no higher than expected by chance for most tools. This observation does not necessarily reflect that there are few consistent genera that differ between obese and non-obese individuals; it could instead simply reflect technical and/or biological factors that differ between the particular datasets we analyzed (Pollock et al., 2018). Despite these complicating factors, it is noteworthy that the MaAsLin2 workflows produced more consistent results than expected based on these datasets.

We believe the above observations regarding DA tools are valuable, but many readers are likely primarily interested in hearing specific recommendations. Indeed, the need for standardized practices in microbiome analysis have recently become better appreciated(Hill, 2020). One goal of our work was to validate the recommendations of another recent DA method evaluation paper, which found that limma voom, corncob, and DESeq2 performed best overall (Calgaro et al., 2020). Based on our results we do not recommend these tools as the sole methods used for data analysis, and instead would suggest using more conservative methods such as ALDEx2 and ANCOM-II. Although these methods have lower statistical power (Calgaro et al., 2020; Hawinkel et al., 2019), we believe this an acceptable trade-off given the higher cost of identifying false positives as differentially abundant. However, MaAsLin2 (particularly with rarefied data) could also be a reasonable choice for users looking for increased statistical power at the potential cost of more false positives. We can clearly recommend that users avoid using edgeR (a tool primarily intended for RNA-seq data) as well as LEfSe for conducting DA testing with 16S data. Users should also be aware that limma voom and the Wilcoxon (CLR) approaches may perform especially poorly on unfiltered data. This is especially true for the Wilcoxon (CLR) approach when read depths greatly differ between groups of interest.

More generally, we recommend that users employ several methods and focus on significant features identified by most tools, while keeping in mind the characteristics of the tools presented within this manuscript. For example, authors may want to present identified taxonomic markers in categories based on the tool characteristics presented within this paper or the number of tools that agree upon its identification. Importantly, applying multiple DA tools to the same dataset should be reported explicitly. Clearly this approach would make results more difficult to biologically interpret, but it would provide a clearer perspective on which differentially abundant features are robust to reasonable changes in the analysis.

A common counterargument to using consensus approaches with DA tools is that there is no assurance that the intersection of the tool outputs is more reliable; it is possible that the tools are simply picking up the same noise as significant. Although we think this is unlikely, in any case running multiple DA tools is still important to give context to reporting significant features. For example, researchers might be using a tool that produces highly non-overlapping sets of significant features compared with other DA approaches. Even if the researchers are confident in their approach, these discrepancies should be made clear when the results are summarized. This is crucial for providing honest insight into how robust specific findings are expected to be across independent studies, which often use different DA approaches.

How and whether to conduct independent filtering of data prior to conducting DA tests are other important open questions regarding microbiome data analysis (Schloss, 2020). Although statistical arguments regarding the validity of independent filtering are beyond the scope of this work, intuitively it is reasonable to exclude features found in only a small number of samples (regardless of which groups those samples are in). The basic reason for this is that otherwise the burden of multiple-test correction becomes so great as to nearly prohibit identifying any differentially abundant features. Despite this drawback, many tools identified large numbers of significant ASVs in the unfiltered data. However, these significant ASVs tended to be more tool-specific in the unfiltered data and there was much more variation in the percentage of significant ASVs across tools. Accordingly, we would suggest performing prevalence filtering (e.g., at 10%) of features prior to DA testing, although we acknowledge that more work is needed to estimate an optimal cut-off rather than just arbitrarily selecting one (McMurdie and Holmes, 2014).

Another common question is whether microbiome data should be rarefied prior to DA testing. It is possible that the question of whether to rarefy data has received disproportionate attention in the microbiome field: there are numerous other factors affecting an analysis pipeline that likely affect results more. Indeed, tests based on rarefied data in our analyses did not perform substantially worse than other methods on average. More specifically, the most consistent inter-tool methods, ANCOM-II and ALDEx2, are based on non-rarefied data, but MaAsLin2 based on rarefied data produced the most consistent results across datasets of the same phenotype. Accordingly, we cannot definitively conclude that rarefying data prior to DA testing is always inadvisable. It should be noted that we are referring only to rarefying in the context of DA testing: whether rarefying is advisable for other analyses, such as prior to computing diversity metrics, is beyond the scope of this work (McMurdie and Holmes, 2014; Weiss et al., 2017).

In conclusion, the high variation in the output of DA tools across numerous 16S rRNA gene sequencing datasets highlights an alarming reproducibility crisis facing microbiome researchers. Unfortunately, this high variation across tools implies that biological interpretations will often drastically differ depending on which DA tool is used. One incomplete solution to this problem would be to normalize the practice of reporting results based on a range of DA tools, which would help ensure that any key conclusions were robust to the researchers’ analysis choices.

## Acknowledgements

We would like to thank everyone who responded to MGIL’s queries on Twitter regarding which differential abundance tools to evaluate. We would also like to thank the authors of all DA tools and datasets used in this study for making their code and data freely available. JTN is funded by both a Nova Scotia Graduate Scholarship and a ResearchNS Scotia Scholars award. GMD was funded by a Canadian Graduate Scholarship (Doctoral) from NSERC. MGIL is funded through a National Sciences and Engineering Research Council (NSERC) Discovery Grant and the Canada Research Chairs program.

