## Supplemental Materials for "Microbiome differential abundance methods produce disturbingly different results across 38 datasets"

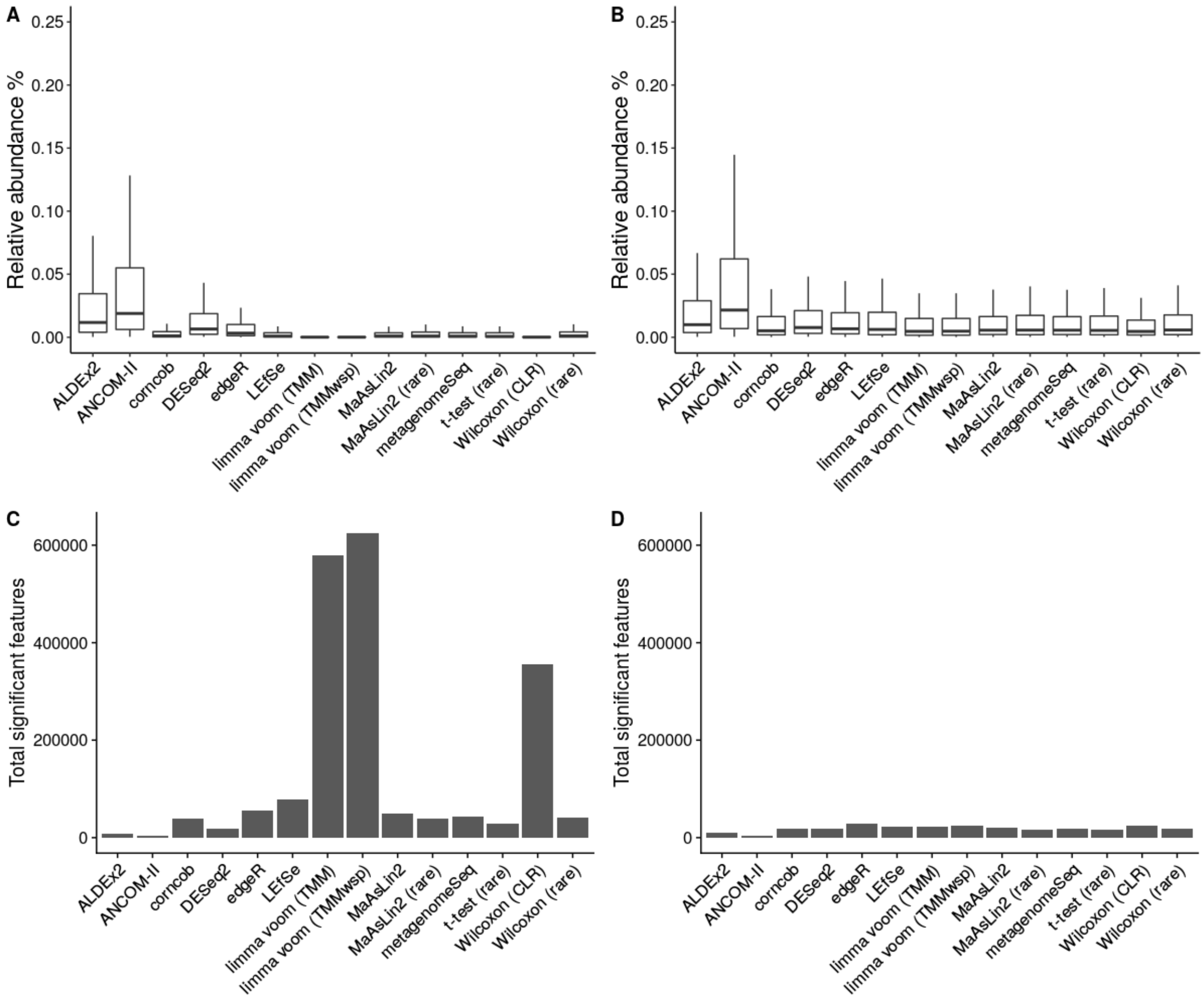


**Supplementary Figure 1: Counts and relative abundances of significant features by tool across all 38 datasets**. (A and B) Boxplots of relative abundance per significant features for the (A) unfiltered and (B) prevalence-filtered approaches. (C and D) Total number of significant features for the (C) unfiltered and (D) prevalence-filtered approaches.

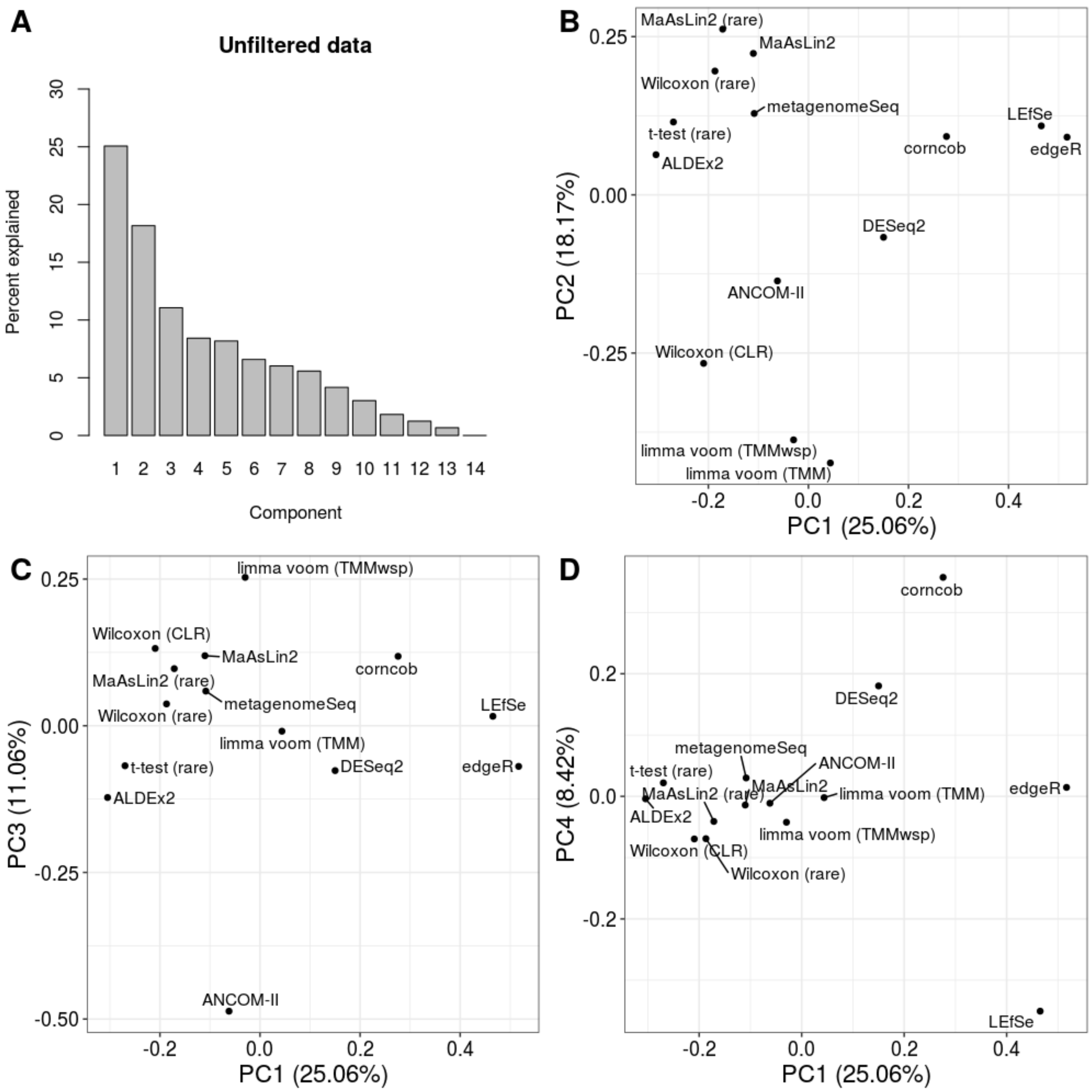


**Supplementary Figure 2: Principal Coordinates Analysis on significant sets of amplicon sequence variants (based on unfiltered data)**. (A) Percentage explained by each component of the Principal Coordinates Analysis (PCoA). (B-D) Two-dimension summaries of PCoA as in Figure 3, but panels C and D visualize components three and four against the first component.

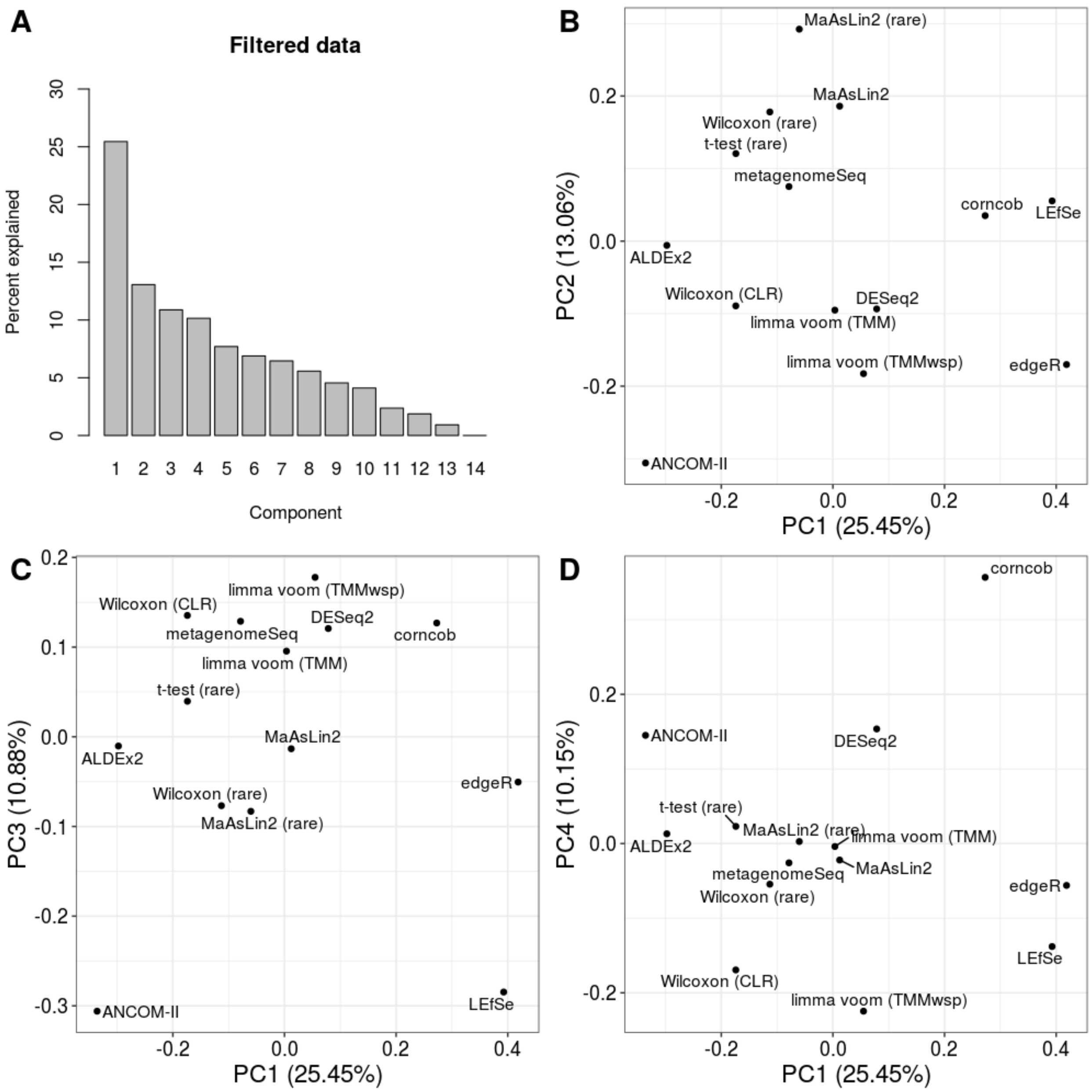


**Supplementary Figure 3: Principal Coordinates Analysis on significant sets of amplicon sequence variants (based on prevalence filtered data)**. (A) Percentage explained by each component of the Principal Coordinates Analysis (PCoA). (B-D) Two-dimensional summaries of PCoA as in Figure 3, but panels C and D visualize components three and four against the first component.

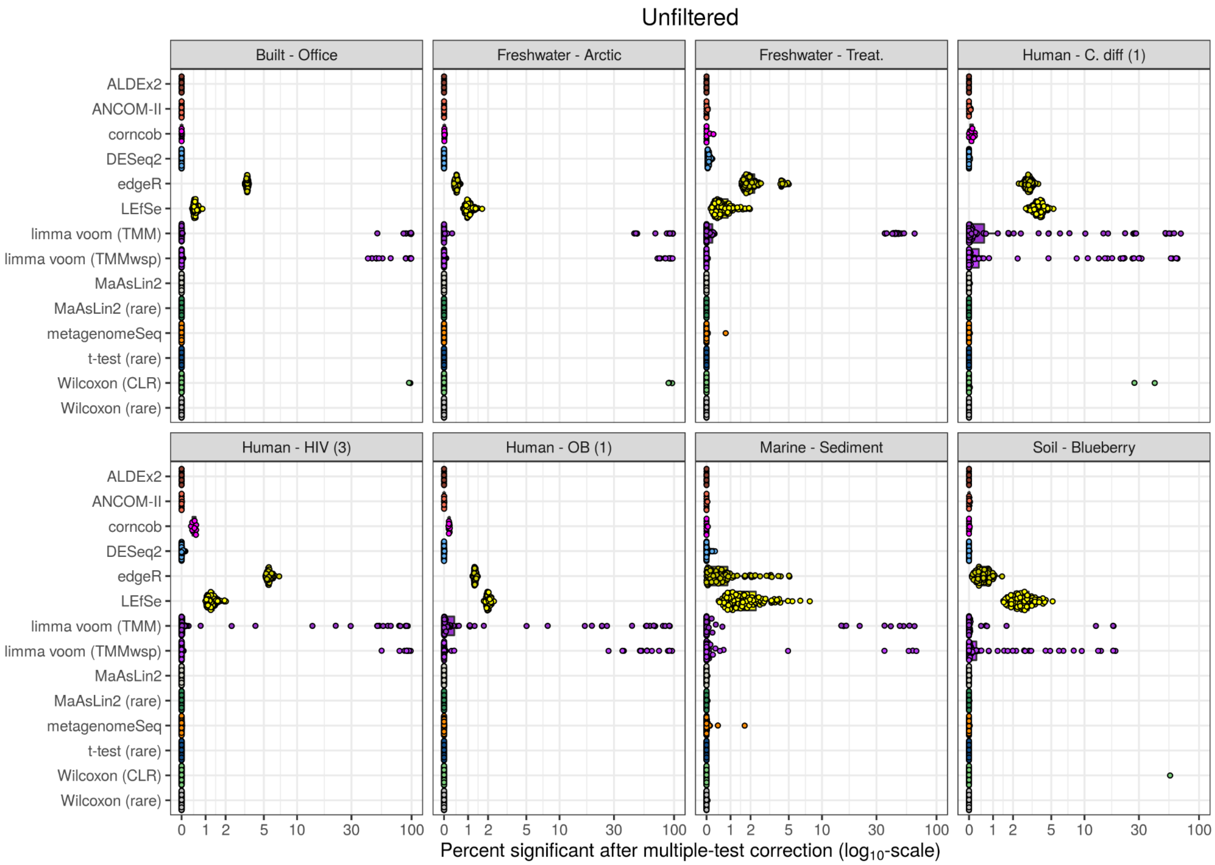


**Supplementary Figure 4: Distribution of false discovery rate simulation replicates for unfiltered data**. The percentage of amplicon sequence variants that are significant after performing Benjamini-Hochberg correction of the P-values (using a cut-off of 0.05) are shown for each separate dataset and tool. These are the raw values that are summarized as averages in Figure 4A. Note that the x-axis is on a $\log_{10}$ scale.

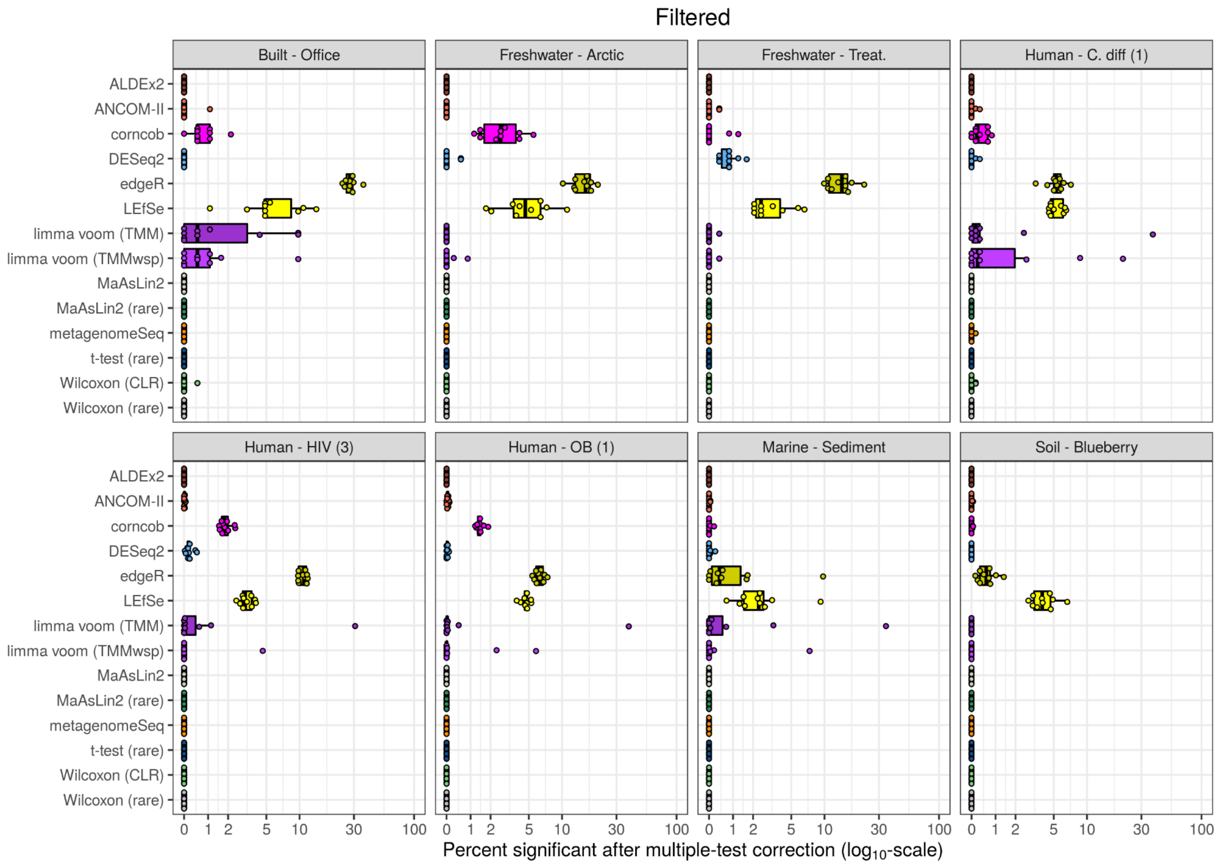


**Supplementary Figure 5: Distribution of false discovery rate simulation replicates for filtered data.** The percentage of amplicon sequence variants that are significant after performing Benjamini-Hochberg correction of the P-values (using a cut-off of 0.05) are shown for each separate dataset and tool. These are the raw values that are summarized as averages in Figure 4B. Note that the x-axis is on a $\log_{10}$ scale.

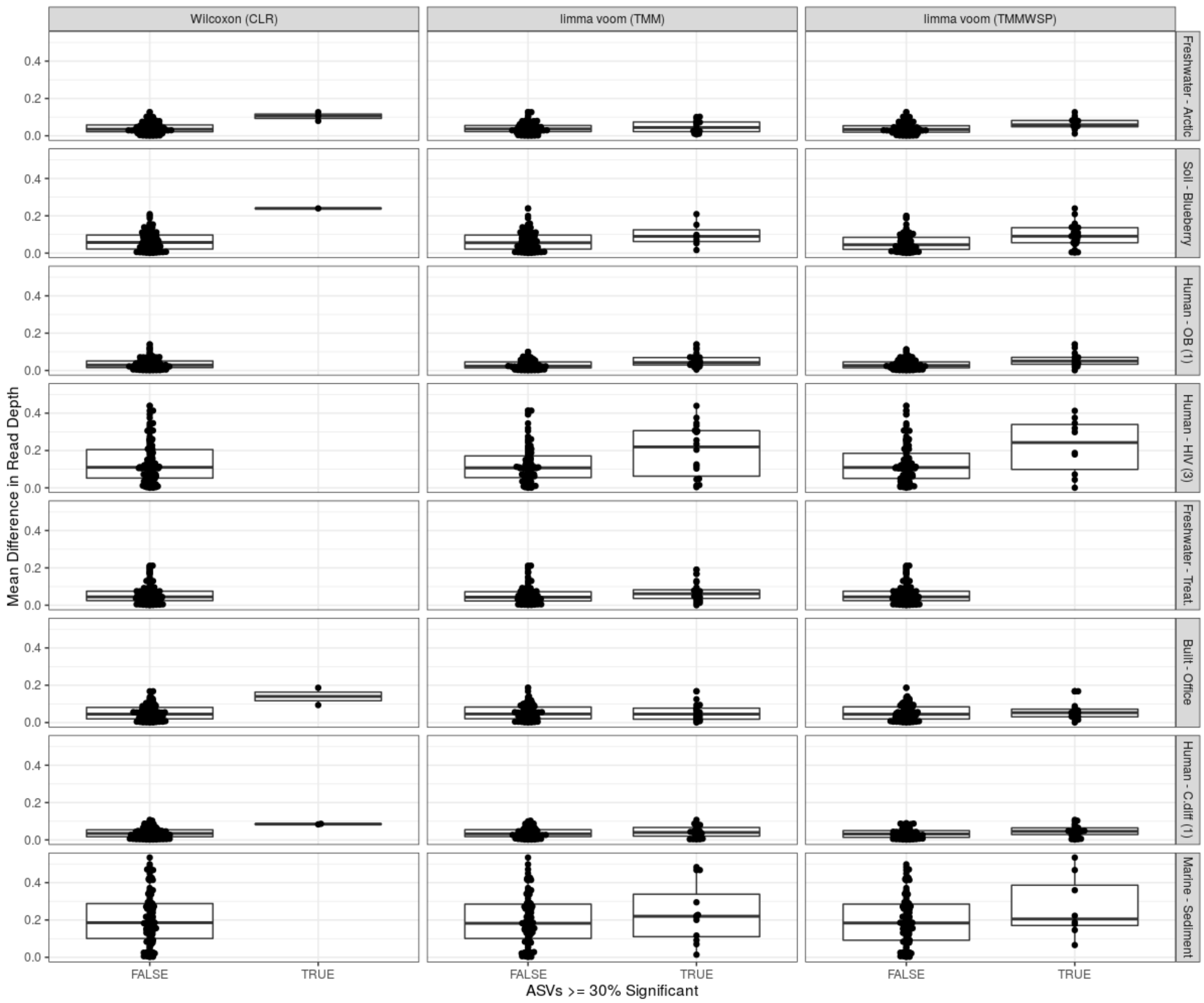


**Supplemental Figure 6: Boxplot comparing mean read depth differences for replicates that resulted in 30% or more ASVs being identified as significant during the false positive analysis.**
For each dataset and replicate in our false positive analysis we checked whether there was a difference in mean read depth between the two tested groups for the Wilcoxon (CLR), limma voom (TMM), and limma voom (TMMWSP). These tools were chosen due to their inconsistent findings across replicates, in some cases identifying greater than 90% of amplicon sequence variants (ASVs) as being differential abundant. The y-axis corresponds to the difference between the group mean read depths normalized by the mean read depth of all samples. On the x-axis, the FALSE category corresponds to replicates where fewer than 30% of ASVs were significant, while TRUE represents replicates where at least 30% or more of the tested ASVs were significant.

**Supplementary Table 1**: Dataset descriptions

| Dataset | Seq. tech. | 16S region | Median depth | Processing Pipeline | Ref. | Raw data provenance |
|---|---|---|---|---|---|---|
| Mouse - Facilities | Miseq | V6-V8 | 6742 | Deblur (ASV) | 10.7717/peerj.5494 | ENA study PRJEB25165 |
| Soil - Blueberry | Miseq | V6-V8 | 11442 | Deblur (ASV) | 10.1094/PBIOMES-03-17-0012-R | SRA study PRJNA389786 |
| Human - ALL | Miseq | V4-V5 | 7171.5 | Deblur (ASV) | 10.3389/fcimb.2019.00028 | ENA study PRJEB29237 |
| Mouse - Exercised | Miseq | V6-V8 | 5212 | Deblur (ASV) | 10.1128/mSystems.00006-17 | ENA study PRJEB18615 |
| Human - CD (1) | Miseq | V6-V8 | 7328.5 | Deblur (ASV) | 10.1186/s40168-018-0398-3 | ENA study PRJEB21933 |
| Built - Office | MiSeq | V4 | 6220 | Deblur (ASV) | 10.1128/mSystems.00022-16 | QIITA study 10423 |
| Human - RA | 454 | V1-V2 | 3216 | USEARCH (de novo OTU) | 10.7554/eLife.01202.001 | SRA study SRP023463 |
| Human - ASD | Miseq | V1-V2 | 4781.5 | USEARCH (de novo OTU) | 10.1371/journal.pone.0137725 | SRA study SRP057700 |
| Human - C. diff (1) | 454 | V3-V5 | 4967 | USEARCH (de novo OTU) | 10.1128/mBio.01021-14 | mothur.org/CDI_MicrobiomeModeling |
| Human - C. diff (2) | 454 | V3-V5 | 2665 | USEARCH (de novo OTU) | 10.1186/2049-2618-1-18 | Duvallet et al. 2017 emailed authors |
| Human - CC (1) | MiSeq | V4 | 11930 | USEARCH (de novo OTU) | 10.1186/s13073-016-0290-3 | SRA study SRP062005 |
| Human - CC (2) | MiSeq | V4 | 120989 | USEARCH (de novo OTU) | 10.15252/msb.20145645 | ENA study PRJEB6070 |
| Human - Inf. | 454 | V3-V5 | 3377 | USEARCH (de novo OTU) | 10.1186/s40168-015-0109-2 | http://dx.doi.org/10.6084/m9.figshare.1447256 |
| Human - HIV (1) | 454 | V3-V5 | 3295 | USEARCH (de novo OTU) | 10.1093/infdis/jiu409 | SRA study SRP039076 |
| Human - HIV (2) | MiSeq | V4 | 3566 | USEARCH (de novo OTU) | 10.1016/j.chom.2013.08.006 | ENA study PRJEB4335 |
| Human - HIV (3) | MiSeq | V3-V4 | 27689.5 | USEARCH (de novo OTU) | 10.1016/j.ebiom.2016.01.032 | SRA study SRP068240 |
| Human - CD (2) | Miseq | V4 | 10014.5 | USEARCH (de novo OTU) | 10.1016/j.chom.2014.02.005 | SRA study SRP040765 |
| Human - IBD | 454 | V3-V5 | 4471 | USEARCH (de novo OTU) | 10.1371/journal.pone.0039242 | Duvallet et al. 2017 emailed authors |
| Human - OB (1) | Miseq | V4 | 27077 | USEARCH (de novo OTU) | 10.1016/j.cell.2014.09.053 | ENA studies PRJEB6702 and PRJEB6705 |
| Human - OB (2) | 454 | V1-V3 | 5038.5 | USEARCH (de novo OTU) | 10.1186/s40168-015-0072-y | SRA study SRP053023 |
| Human - OB (3) | 454 | V2 | 2831 | USEARCH (de novo OTU) | 10.1038/nature07540 | https://gordonlab.wustl.edu/NatureTwins_2008/TurnbaughNature_11_30_08.html |
| Human - OB (4) | 454 | V4 | 9778 | USEARCH (de novo OTU) | 10.1002/hep.26093 | MG-RAST, study mgp1195 |
| Human - Par. | 454 | V1-V3 | 2476.5 | USEARCH (de novo OTU) | 10.1002/mds.26069 | ENA study PRJEB4927 |
| Human - T1D (1) | MiSeq | V4 | 9885 | USEARCH (de novo OTU) | 10.2337/db14-1847 | Duvallet et al. 2017 emailed authors |
| Human - T1D (2) | 454 | V4 | 4702.5 | USEARCH (de novo OTU) | 10.1038/srep03814 | Duvallet et al. 2017 emailed authors |
| WWSR - Continents | MiSeq | V4 | 35206 | Mothur (do novo OTU) | 10.1038/s41564-019-0426-5 | SRA study PRJNA509305 |
| WWSR - Temp. | MiSeq | V4 | 36791 | Mothur (do novo OTU) | 10.1038/s41564-019-0426-5 | SRA study PRJNA509305 |
| Marine - Sediment | MiSeq | V3-V4 | 25624 | Deblur (ASV) | 10.1021/acs.est.5b01093 | SRA study SRA233339 |
| Marine - Plastic (1) | MiSeq | V4-V5 | 75591 | Deblur (ASV) | 10.1016/j.envpol.2018.07.023 | VAMPS portal LQM_MPLA_Bv4v5 |
| Marine - Plastic (2) | MiSeq | V4 | 16664 | Deblur (ASV) | 10.1086/693012 | SRA study SRP077684 |
| Marine - Plastic (3) | MiSeq | V4 | 34990 | Deblur (ASV) | 10.3389/fmicb.2019.01665 | SRA study PRJNA506548 |
| River - Plastic | MiSeq | V4 | 18942 | Deblur (ASV) | 10.1002/ecs2.1556 | SRA study SRP065321 |
| Marine - Plastic (4) | MiSeq | V4 | 4545 | Deblur (ASV) | 10.1371/journal.pone.0159289 | SRA study PRJNA283545 |
| Marine - Plastic (5) | MiSeq | V1-V3 | 10851 | Deblur (ASV) | 10.1016/j.scitotenv.2019.135790 | SRA study PRJNA602877 |
| Soil - Arctic | MiSeq | V4 | 22472 | Deblur (ASV) | Not found | ENA study ERP111883 |
| Soil - Fires | HiSeq | V4 | 71394 | Deblur (ASV) | Not found | ENA study ERP016543 |
| Freshwater - Arctic | HiSeq | V4 | 3272 | Deblur (ASV) | Not found | ENA study ERP017459 |
| Freshwater - Treat. | MiSeq | V4 | 65455 | Mothur (do novo OTU) | 10.1371/journal. pone.0141087 | QIITA study 10251 |
| dia_schneider* | MiSeq | V3-V4 | 38192 | USEARCH (de novo OTU) | 10.1038/sdata.2017.152 | SRA study PRJNA353353065 |
| GEMS1* | 454 | V1-V2 | 3536 | DADA2 (ASV) | 10.1186/gb-2014-16-6-r76 | https://microbiomedb.org/mbio/app/record/dataset/DS_bb7b589593 |

**Used solely for consistency analyses with diarrhea datasets*

**Supplementary Table 2: Consistency of significant genera calls across obesity datasets**

| Tool | No. sig. genera | Max overlap | Mean exp. | Mean obs. | Fold diff. | D | P |
|---|---|---|---|---|---|---|---|
| MaAsLin2 (rare) | 8 | 2 | 1.018 | 1.25 | 1.228 | 0.232 | 0.006 |
| MaAsLin2 | 20 | 3 | 1.071 | 1.25 | 1.167 | 0.13 | 0.029 |
| t-test (rare) | 6 | 2 | 1.011 | 1.167 | 1.154 | 0.156 | 0.038 |
| ALDEx2 | 24 | 3 | 1.063 | 1.167 | 1.098 | 0.063 | 0.197 |
| limma voom (TMMwsp) | 34 | 3 | 1.127 | 1.235 | 1.096 | 0.084 | 0.151 |
| corncob | 34 | 3 | 1.136 | 1.206 | 1.062 | 0.051 | 0.321 |
| limma voom (TMM) | 44 | 3 | 1.171 | 1.205 | 1.029 | 0.02 | 0.75 |
| Wilcoxon (rare) | 22 | 2 | 1.06 | 1.091 | 1.029 | 0.031 | 0.541 |
| edgeR | 68 | 3 | 1.327 | 1.338 | 1.008 | 0.019 | 0.849 |
| DESeq2 | 40 | 2 | 1.123 | 1.125 | 1.002 | 0.005 | 0.923 |
| Wilcoxon (CLR) | 47 | 3 | 1.17 | 1.17 | 1 | 0.013 | 0.818 |
| metagenomeSeq | 4 | 1 | 1.004 | 1 | 0.996 | 0.004 | 0.528 |
| LEfSe | 53 | 3 | 1.203 | 1.189 | 0.988 | 0.039 | 0.437 |
| ANCOM-II | 8 | 1 | 1.017 | 1 | 0.983 | 0.017 | 0.559 |

**Column descriptions:**
**No. sig. genera**: Number of genera significant in at least one dataset
**Max overlap**: Max number of datasets where a genus was called by significant by this tool
**Mean exp.**: Mean number of datasets that each genus is expected to be significant in (of the genera that are significant at least once)
**Mean obs.**: Mean number of datasets that each genus was observed to be significant in (of the genera that are significant at least once)
**Fold diff.**: Fold difference of mean observed over mean expected number of times significant genera are found across multiple datasets
**D**: KS-test statistic
